# RNA multiscale simulations as an interplay of electrostatic, mechanical properties, and structures inside viruses

**DOI:** 10.1101/2023.03.30.535003

**Authors:** Sergio Cruz-León, Salvatore Assenza, Simón Poblete, Horacio V. Guzman

## Abstract

Multiscale simulations have broadened our understanding of RNA structure and function. Various methodologies have enabled the quantification of electrostatic and mechanical interactions of RNA at the nanometer scale. Atom-by-atom simulations, coarse-grained strategies, and continuum models of RNA and its environment provide physical insight and allow to interpret diverse experiments in a systematic way. In this chapter, we present and discuss recent advances in a set of methods to study nucleic acids at different scales. In particular, we introduce details of their parameterization, recent applications, and current limitations. We discuss the interaction of the proteinacous virus capsid, RNA with substrates, compare the properties of RNA and DNA and their interaction with the environment, and analyze the application of these methods to reconstruct the structure of the virus genome structure. Finally, the last lines are dedicated to future developments and challenges ahead.

## 1 Introduction

RNA research is thriving in physical virology. Several packing mechanisms of a few counted viruses and emerging functions of RNA structure inside virus shells have been already elucidated [1]. However, the assembly of RNA viruses is complex and involves the interaction between the genome and the capsid proteins. RNA viruses typically have genomes of several thousand nucleobases, highly charged, structurally complex, and densely packed within the virion. Understanding the physics of viral RNA requires addressing diverse phenomena that occur on different temporal and spatial scales. In this chapter, we will show recent advances in the multiscale modeling of RNA. We will start by discussing the importance of electrostatic effects (section 1). On the one hand, we will use continuum theories to describe the interaction of RNA with surfaces and the impact of nucleic acid secondary structure. On the other hand, we describe electrostatics from the detailed insight of atomistic simulations. The simulations provide access to the thermodynamics and kinetics that govern interactions between RNA and its environment. In section 3, we focus on the elastic properties of nucleic acids. We review the experimental and computational methods to determine these properties and show advances in all-atom and coarsegrained models for an accurate description of the mechanical complexity of nucleic acids. In section 4, we offer algorithms that allow the assembly of the RNA tertiary structure using fragments. Finally, we discuss current challenges and the future role of realistic multiscale modeling in advancing our understanding of RNA biophysics.

## 2 The importance of RNA electrostatic interactions

Electrostatic interactions are often prescribed as one of the generators, or the most important one in virus capsids [2], as well as full virion assembly [3]. The precise number of charges carried by the proteins and genome of virions depend strongly on the solution conditions and structural distribution of dissociable groups on the regions close to the solvent accessible surface. A major part of the volume of the virion is compose of DNA or RNA, which are macro-ions that carry massive amounts of charge, one phosphate group (negative charge) per nucleotide. Consequently, nucleic acids form an ionic atmosphere composed of excess of cations and depletion of anions around that also screen the electrostatic interactions. Without the ionic atmosphere, the confinement inside the viral capsid, folding, and function of nucleic acids would involve crossing enormous electrostatic barriers [4]. At physiological conditions, the ionic atmosphere for RNA viruses is commonly located close to the proteinaceous capsid, which also converges with the concept of neutral total charge inside the capsid required for spontaneous assembly [5]. The ionic atmosphere around nucleic acids is divided into diffusive and associated ions [6]. On the one hand, diffusive ions refer to a significant accumulation of mobile cations (and depletion of anions) around the polyelectrolyte determined largely by long range electrostatics. The diffusive layer contains the majority of the interacting ions, contributes significantly to the thermodynamic stability of DNA and RNA secondary structures [6, 7], and regulates the adsorption of RNA to substrates [8] (section 2.1). On the other hand, the associated ions bind in specific sites of nucleic acids [7, 9]. Their effects strongly depend on the ion type and its understanding requires an atomistic level of description (section 2.2.1). Associated ions induce conformational changes and affect the stability, melting temperature, folding kinetics, and biological activity of DNA and RNA (see reference [9] for an overview).

Furthermore, ions screen electrostatics, stabilize DNA and RNA and can be responsible for large conformational changes [10], such as the compactness of the genome inside a virus capsid. They can induce bending [11, 12], over- and underwinding [13, 14, 15], stabilize tertiary structures and assist the catalytic activity of RNAs [7, 6, 16, 17]. Therefore, to understand nucleic acids, sophisticated knowledge and modeling of the ions around them is crucial [4]. From the protein side, there is also crucial to understand the molecular and even atomistic identity [18], as well as the charged states [1].

Given the scales of virus genome, providing insight into the interpretation of electrostatics phenomenology requires an integrative multiscale approach. Atomistic simulations can provide a high level of detail but are limited to short time- and length scales. On the other hand, coarse-grained (CG) simulations or continuous models allow us to access larger systems, but their insight is finite. For example, the SARS-CoV-2 reach 30 kbp and several millions of residues [19, 20]. The sampling of a system like SARS-CoV-2 in all-atom simulations is still computationally prohibitive. Here, coarse-grained methods have the opportunity of mediating between all-atom and continuum approaches [21, 22]. However, an integrative CG model for both proteins and RNA with tuned electrostatics is still a challenging topic [8, 23, 24, 25]. From a continuum Poisson-Boltzmann approach the ionic clouds surrounding RNA molecules have been also observed [26, 9]. Although continuous models may overlook associative ions at atomistic scale, they still provides *grosso modo* insight. Understanding viral electrostatics requires feedback loops between different modeling methods at different scales, in order to complement each other.

### 2.1 Capsid proteins and RNA-substrate interactions with Coarse-Grained and Continuum models

Recent research results, show the electrostatic interactions of a nanoprobe with the proteins inside and at the surface of a Zika Virus capsid described as a function of the distance, molecular charge identity and variations of ionic strength [18]. RNA as flexible molecules go through structural changes to adsorb onto substrates. The problem of how the secondary structure regulates the adsorption mechanisms of RNA is unsolved from a generalized perspective. Here, we take a major step in its solution by creating a multiscale method, based on electrostatic Debye-Hückel theory, to calculate the adsorption free energies between archetypical RNA fragments and structureless flat substrates. The resulting model, accounting with existing secondary structures obtained from chemical probing experiments, predicts the adsorption free energies and connects to RNA 3D structure [27, 28]. The latter modulate the interaction regimes, for more or less efficient adsorption. This works sets the stage for tackling RNA interactions to proteinaceous substrates, such as virus capsids and the effects those substrates may have on the elemental RNA secondary structure and vice versa.

#### 2.1.1 Localized proteinaceous capsid electrostatic interactions in variable salinity environments

A continuum description of electrostatic interactions is crucial for understanding function and phenomenology in biological systems. In particular, the interactions generated at the atomistic and molecular resolutions are of predominant interests for non-enveloped and enveloped viruses [29, 30, 31, 32, 33, 34, 1, 35]. For a few decades, the physical virology community has employed diverse experimental techniques, like force microscopy, to learn more about the mechanical and electrostatic properties of biomacromolecular systems [36, 37, 38, 39, 40, 41, 42, 43, 44, 45, 46]. However, the amount of viruses that are electrostatically characterized at the nanoscale (*i.e.* few nanometers) are scarce, where we can highlight the following pioneering results [38, 18]. The most recent computational model [18] tackles the electrostatic force arising from the interaction between a nanoprobe and the virus capsid via Poisson-Boltzmann calculations.

There are several ways to calculate the electrostatic forces using PoissonBoltzmann [47]. Here, the total force **F** is decomposed as,

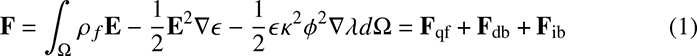

where *ρ _f_* is the charge distribution in the solute, **E** the electric field, and *λ* a unit step function that is 1 in Ω*_w_* (the region containing the solvent- water) and 0 elsewhere, *ϵ* is the dielectric constant, *κ* is th inverse of the Debye length and *Φ* as the electrostatic potential. The first term **F**_qf_ corresponds to the force on the solute’s charges, then **F**_db_ corresponds to the pressure due to the dielectric jump on the interface, and the final term is **F**_ib_ arises from the sudden jump in ionic concentration on the molecular surface. More details on the derivation of equations and definition of variables are contained elsewhere [18].

Fig. 1 shows the interaction force (force magnitude normalized by the total magnitude **F** _qf_), indicating that the atoms located closer to the tip are also concentrating most of the contribution to the total force (blue atoms in Fig. 1(C) and (D)). Note that the nanoprobe approaches from the right end in the x-axis. From the experimental AFM viewpoint, the dependence on the distance is a well-known observable of the sensed molecular interactions. In order to cover this aspect from the theoretical perspective, we analyzed the atoms corresponding to each amino acid based on 5 Å thick slices (normal to the *x*-direction), which resembles the tip-capsid distance in a similar form as AFM ’force tomography’ or subsurface imaging [48]. Note that Fig. 1(C) and (D) differ only in the amount of ionic concentration in the solvent, namely, 20mM driving stronger electrostatic interactions and 150mM, which is rather screening most of the ions. The contribution of the atoms from specific amino acids is reflected in the force tomography profiles in Figs. 1(A) and 1(B). In Fig. 1(A), we identify 5 amino acids that are more sensitive to the nanoprobe, namely, LYS, ASP, GLU, ARG and ILE. Those amino acids have different net charges depending on the pH of the medium where they are located, whereby Fig. 1(A) shows that for LYS and ARG the interactions are attractive, while for both ASP and GLU are of repulsive nature. Nonetheless, the case of ILE are much more sensitive to the slice where they are located. At pH 7, ILE is an uncharged residue, however if they are sliced by the tomography-type analysis (the whole amino acid is distributed across several slices), the sensed polarity by the nanoprobe depends on the exact charges of the atoms of the specific slice (Fig. 1(A)). This explains the observed transitions from repulsive to attractive interactions and vice versa. Interestingly, the same quantification was performed for the higher salt concentration force, namely 150mM (Fig. 1(B)), here the amino acids with higher contribution to the force have changed drastically in order and also in species, as it is the case for LYS, THR and SER.

**Fig. 1.**
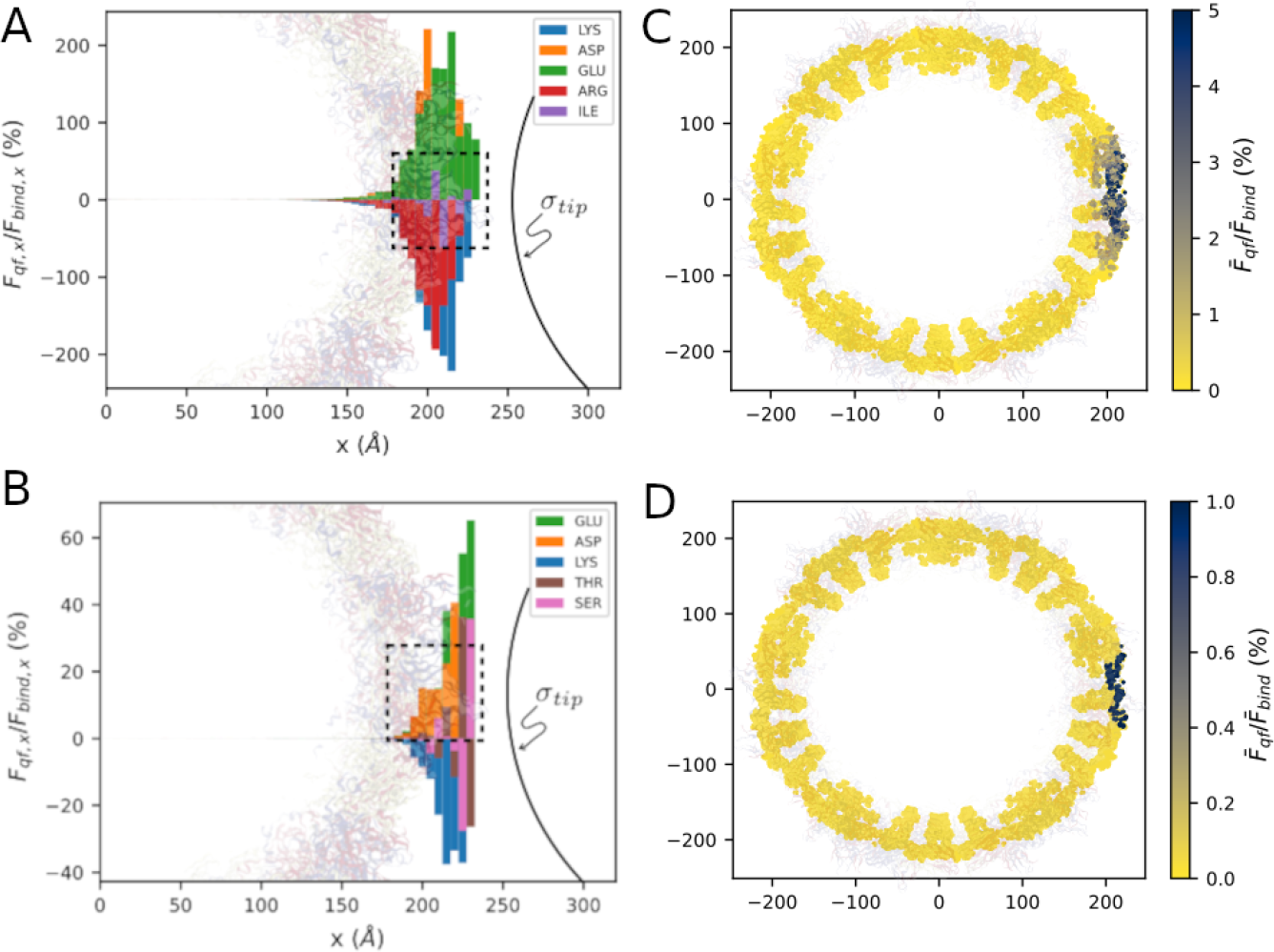
(A) The top left panel shows the x-component of the interaction force (normalized) on atoms that belong to the 5 amino acids that contribute the most to the force (LYS, ASP, GLU, ARG, and ILE, in that order) on 5 Å-thick slices. As reference, we show the Zika capsid structure in the background, and the AFM tip, which is placed 2 Å away. (B) A similar plot as (A) with a remarking difference, now the ionic concentration is 150mM and hence the indentity of the top 5 amino acids drastically changes in order and species (GLU, ASP, LYS, THR, and SER, in that order). (C) Relative magnitude of **F** *_q,f_* on atoms located in a 5 Å slice centered at the *z* = 0 Å plane at ionic concentration is 20mM and (D) 150mM. Figure reproduced from [18] with licence CC-BY 3.0.

In summary of this subsection, we showed that a simple electrostatic model based on the Poisson-Boltzmann equation provide useful insights of fine-grain mechanisms in the interaction between an AFM tip and a virus capsid. In particular, we demonstrate that the electrostatic force originated from a proteinaceous capsid and detected by a nanoprobe depends on both the precise distribution and identity of charges inside the amino acids and the tip-capsid distance. The research highlight the strong locality of the total interaction force with the amino acids as it approaches the nanoprobe. This results also suggest that the protein shell may shield the RNA genome inside the capsid at higher ionic concentrations. However, further verification, from the side of the RNA molecule inside are currently in discussion [1]. As a consequence, the electrostatic component of the force requires higher levels of resolution when modeling and interpreting heterogeneous surfaces, like the ones of virus capsids.

#### 2.1.2 RNA adsorption onto flat substrates

Molecular simulations have offered important insights into the adsorption of semiflexible macromolecules to a molecular substrate [49, 50, 51, 52], while the adsorption of macromolecules with either double-stranded or quenched internal structure onto a molecular substrate remains much less understood. The latter problem is particularly relevant in the context of RNA-virus assembly phenomena [53, 54, 3, 55, 56, 57, 22, 1], where the soft and malleable RNA structure can respond to the adsorption process.

To investigate the general role of RNA secondary structure in adsorption processes, we model the substrate as a flat, featureless surface. Excluding topographical and molecular features of the substrate, such an scenario allows us to isolate the influence of the RNA secondary structure in its adsorption. The employed RNA model for unstructured fragments is similar to a single chain bead-spring polymer while the structured RNA uses a tractable CG scheme [58]. We model the attraction of the RNA phosphate groups to the adsorbing substrate by a Debye-Hückel-like interaction potential, which can be rationalized as stemming from the electrostatic interactions between the dissociated RNA phosphates and the substrate charges [59]. Combined with a generic short-range repulsive term [8], the surface potential acting on the RNA then assumes the form,

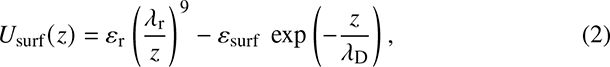

where *z* is the distance to the surface, *ε*_r_ = 1 *k*_B_*T*, *λ*_r_ = 0.1 nm is the distance of activation of the repulsive Lennard-Jones term (well below the size of the RNA phosphates ≈ 0.3 nm, as defined in our model) and *λ*_D_ = 1 nm, considering RNA under typical physiological conditions as described in Refs. [60, 61]. The strength of the attractive potential, *ε*_surf_, is varied in the range between 0.44 *k*_B_*T* and 1.78 *k*_B_*T*, which are also observed experimentally for nucleic acid packaging [62].

Figure 2(A) shows the adsorption free energy *F*_m_ (minimum of the PMF) as a function of the surface attraction strength *ε*_surf_, for both structured (wiss) and unstructured (noss) RNA systems. The tackled fragment is system 1 (S1), a nucleotide RNA hairpin, with the following secondary structure *((.(((((.))))).))*.

**Fig. 2.**
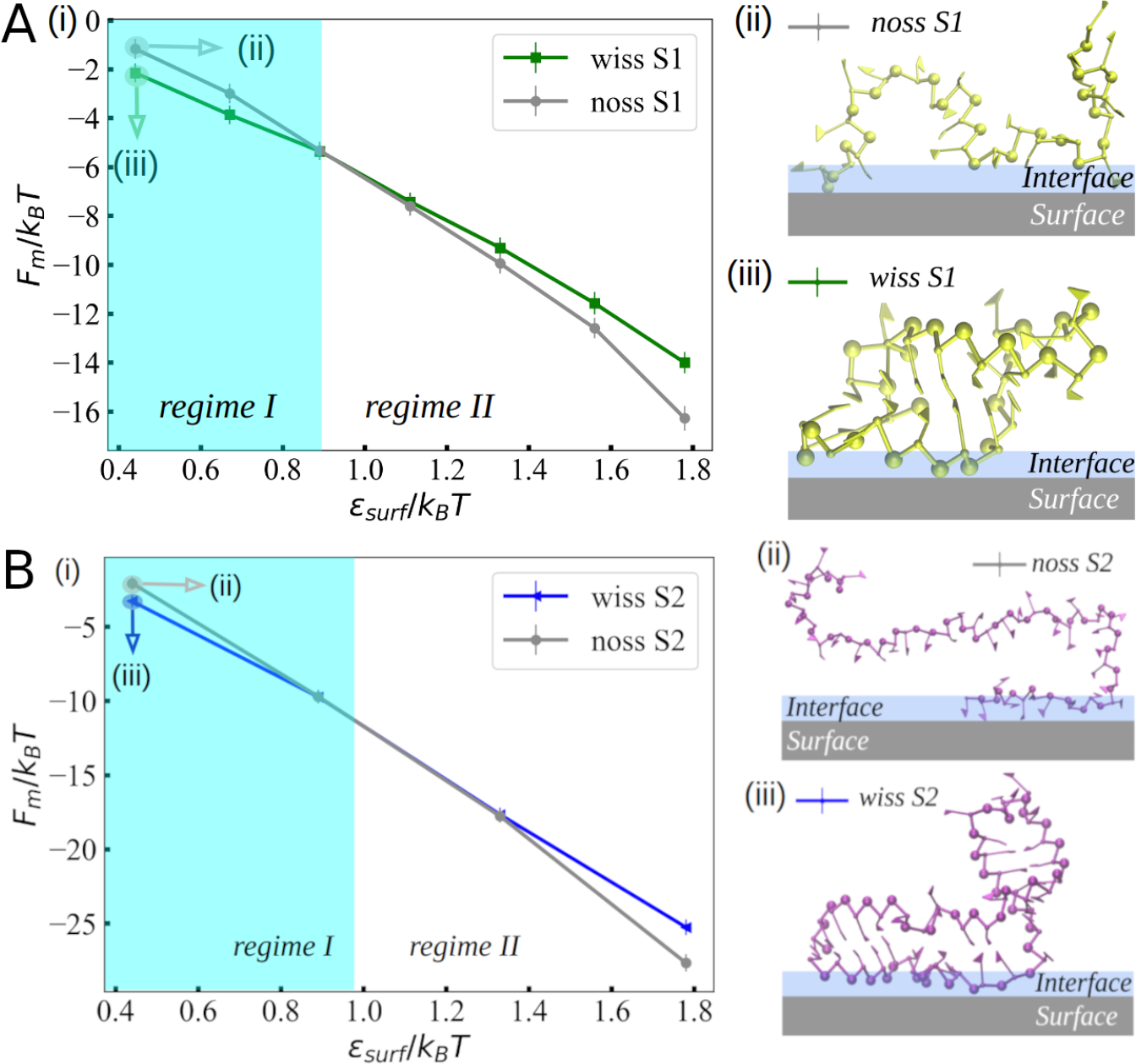
(A) Adsorption free energy as a function of the substrate attraction strength, *ε*_surf_/*k*_B_*T*, for RNA fragments S1-wiss and S1-noss. For a weak attraction of *ε*_surf_/*k*_B_*T* = 0.44, we show simulation snapshots of the (ii) unstructured and (iii) structured fragments. The illustrated interface layer is a schematic definition of a distance slightly thicker than the diameter of phosphates. (B) Adsorption free energy *F_m_* as a function of the substrate attraction strength *ε*_surf_ for RNA fragments S2-wiss and S2-noss. Snapshots of (ii) unstructured and (iii) structured fragments at *ε*_surf_ = 0.44 *k*_B_*T*. Figure reproduced from [8] with licence CC-BY 4.0.

Two regimes are identifiable: the first for *ε*_surf_ *<* 0.89*k*_B_*T*, where the fragment with secondary structure adsorbs more strongly than the unstructured one (regime I), while for *ε*_surf_ *>* 0.89*k*_B_*T*, the unstructured fragment is the one exhibiting a stronger adsorption free energy (regime II). An intriguing question is how general is the existence of the two adsorption regimes. To provide a rough answer, we performed simulations using a different type of attractive potentials [8]. Moreover, we studied another archetypical RNA fragment of the STMV shape secondary structures [63]. System 2 (S2) contains 40 nucleotides and a slightly lower base-pair fraction *(((((…((.(((((……))))).)).)))))*, namely 63% (S1), 60% (S2). This brings into play the shifting of the cross-over point to the right, which remarks the sensitivity of the molecule during the adsorption phenomena (as shown in Fig. 2(B)).

Our results, which indicate a selectivity in adsorption between singleand doublestranded regions of RNA, underline the importance of RNA structure in regulating its adsorption to various substrates. We expect that the selective adsorption of one RNA structure over the other could be experimentally controlled by tuning the interaction strength, for instance, by changing the salt concentration, salt type or pH. Moreover, the lower interaction free energies of unstructured RNAs compared to structured, double-stranded ones at high attractions suggest that possibly highly attractive surfaces may promote the unfolding of a double-stranded RNA structure.

### 2.2 RNA electrostatics at the atomic level: The importance of the details

At the molecular level, the interaction between RNA and the surrounding ions and water completely determines the electrostatics. All-atom molecular dynamics simulation (MD) is powerful computational technique that allows investigating physical systems with molecular resolution. MD is a “computational microscope” [64], and it is well suited to study RNA electrostatics because MD can account for the identity of ions, the critical role of water, the internal flexibility of nucleic acids, and the effect of the many-body problem with the required spatial and time resolution. MD has been successfully used in modeling many biological systems atom-by-atom. Recently and thanks to the effort of the community, MD models has proven also powerful in quantitatively reproducing the conformations dynamics and mechanical response of DNA and RNA [65, 66, 67, 15, 68, 69, 70, 71] (see section 3). This section shows how MD provides molecular insight into the interactions of RNA with ions. This atomic view is essential to understand complex processes such as RNA packaging into viruses because ions can modify the physical and chemical properties of RNA and modulate its interaction with other molecules.

#### 2.2.1 Ion-specific effects in RNA systems

Multiple physical and chemical properties of RNA systems depend on the ion type present in the solution. In this section, we study the thermodynamics and kinetics of ion binding in specific sites of nucleic acids [7, 9] using an atomistic description. We used MD simulations using recently improved models [72, 73, 71, 65, 74] combined with enhanced sampling techniques to investigate the interactions of eight mono- and divalent cations with the main binding sites of RNA [9]. The main binding sites on RNA correspond to the backbone (non-bridging oxygen atoms of the phosphate groups: atoms O1P and O2P) and nucleobase (atoms N7 and O6) binding sites. Using a dinucleotide (snapshots in Figure 3), we investigated the binding mechanism, affinities, and exchange rates of Li^+^, Na^+^, K^+^, Cs^+^, Mg^2+^, Ca^2+^, Sr^2+^, and Ba^2+^ to understand the molecular origin of ion-specific effects on RNA.

**Fig. 3.**
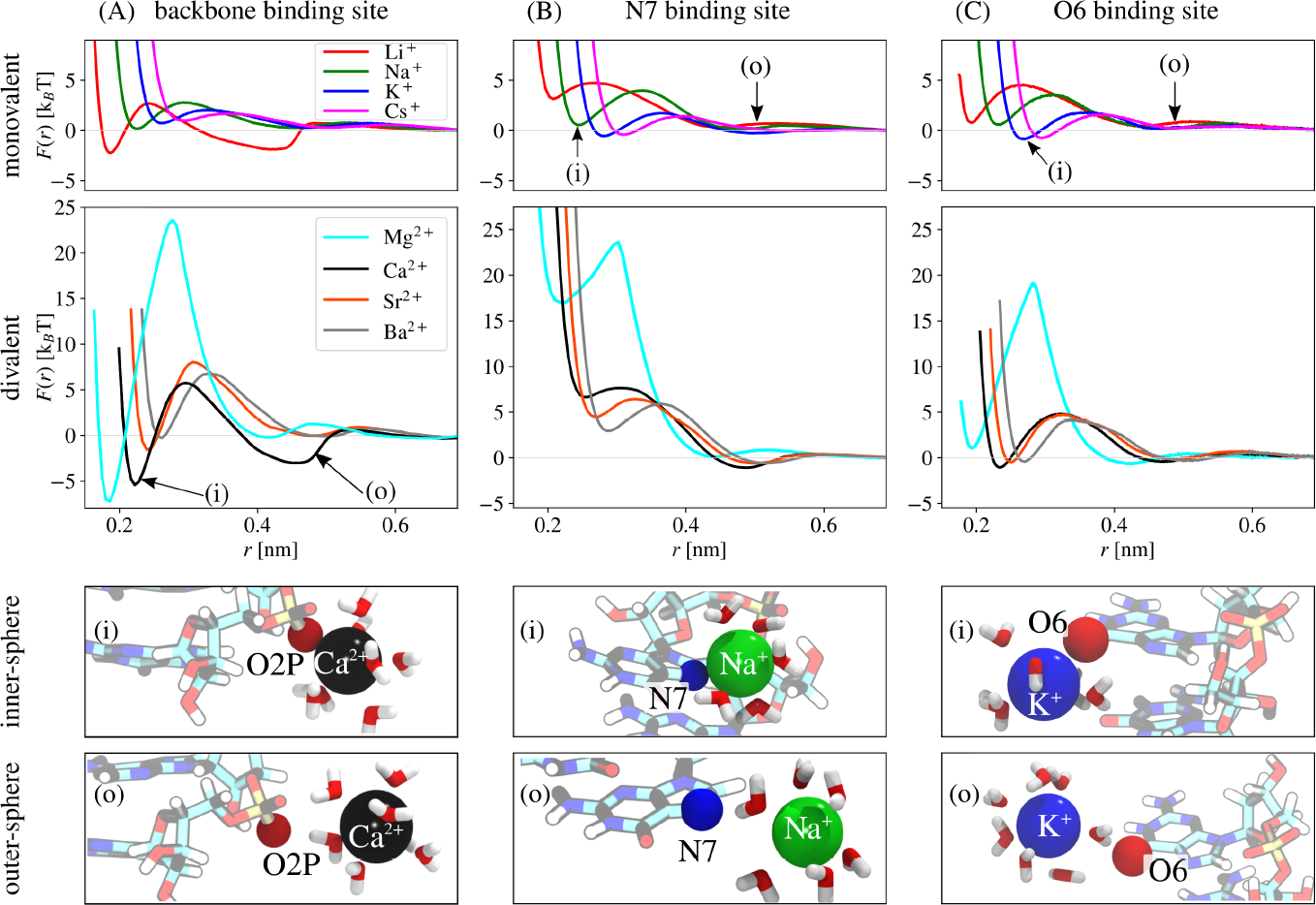
Free energy profiles *F* (*r*) for the binding of metal cations as a function of the distance to the O2P atom in the backbone binding site (A), and to the N7 (B) and the O6 binding site in the nucleobase (C). Simulation snapshots for inner-sphere (i) and outer-sphere (o) conformation at the different ion binding sites (bottom). The arrows indicate at which separation the snapshots were taken. For clarity, only water molecules in the first hydration shell of the cation are shown. Figure reproduced from [9] with licence CC-BY 4.0.

Fig. 3 shows the free energy profiles *F* (*r*) for the metal cations as a function of the distance to the backbone (Fig. 3A) and nucleobase binding sites (Figs. 3B and 3C)^1^. In all cases, the free energy profiles exhibit two stable states separated by an energetic barrier that shows a common binding mechanism. The first minimum is an inner-sphere interaction where a partially dehydrated cation is in direct contact with the binding site (snapshots (i)). The second minimum is an outer-shell contact, where a fully hydrated ion interacts with the RNA site mediated by a water molecule (snapshots (o)). Therefore, ion-water interactions strongly influence ion binding to RNA.

The free energy profiles show opposing trends for binding affinities. First, at the backbone binding site, the first minimum in F(r) gets deeper with decreasing ion diameter (Fig. 3A). The depth of the first minimum indicates that ions with higher charge density (such as Li^+^ or Mg^2+^) interact stronger with the non-bridging oxygen atoms of the RNA backbone (O1P and O2P) and, consequently, have a higher binding affinity. This effect help to explaining changes in free energies [16, 71, 73] and nicely correlates with the ability of ions like Ca^2+^ and Mg^2+^ to stabilize RNA structures [75, 7, 9] The trend is completely reversed for the nucleobase binding site N7. At the N7 site, ions with low charge density are preferred, which provides further evidence for the discussion on the binding of Mg^2+^ in nucleic acids [76].

Finally, the binding kinetics is also affected by ion-type and RNA binding site. For example, at the backbone binding site, the energetic barrier separating the two stable states increases with decreasing ion diameter, therefore slowing exchange kinetics. We determined that ion-binding lifetimes span more than five orders of magnitude, from picoseconds for Cs^+^ up to hundreds of nanoseconds for Ca^2+^ [9, 71]. The vast changes ion-binding lifetimes helps to understand .

Overall, the results from the dinucleotide reveal notorious ion-specific effects and a high selectivity of RNA binding sites. The site-specific affinities and the vast time-scale involved provides a microscopic explanation of the ion-specific effects observed in multiple macroscopic RNA properties. For example, the results from Fig 3 help us to understand the central role of ion type in RNA folding stability and kinetics. RNA stability is largely determined by the binding affinity of the ions. On the other hand, RNA folding times depend on binding kinetics, and therefore is affected by lifetimes on ion-binding. However, although conceptually useful, the dinucleotide results are limited when one tries to understand the complex structural landscape of nucleic acids. Consequently, in the following section, we use atomistic simulations to quantify the effects of ions in the more structurally complex nucleic acids.

#### 2.2.2 Electrostatics around DNA and RNA duplexes

Ion distributions and nucleic acid structure are intertwined: ion distributions modify the nucleic acid structure [14, 13, 15]; conversely, nucleic acid structure modifies ion distributions [77]. In this section, we use MD simulations to show how the ion distributions, and consequently the electrostatics around dsDNA and dsRNA, change depending on ion type and the nucleic acid type. We focus on duplexes because they are the main form of DNA and the most common secondary fragment for RNA. For example, the crystal structure of the Satellite Tobacco Mosaic Virus reveals up to 60% of the genome folded into helices [78].

Fig. 4 shows the ion distributions obtained from extensive unrestrained simulations of 33-base pairs (bp) DNA and 33-bp RNA duplexes in their B-helix and Ahelix topologies, respectively (see details in Ref. [77]). To allow the identification and comparison n specific nucleic acid volumes such as the major groove, minor groove, and the non-bridging oxygen atoms in the phosphate groups along the back-bone, we used the curvilinear helicoidal coordinate system introduced by Laverly et al., [79] (Fig. 4B,D).

**Fig. 4.**
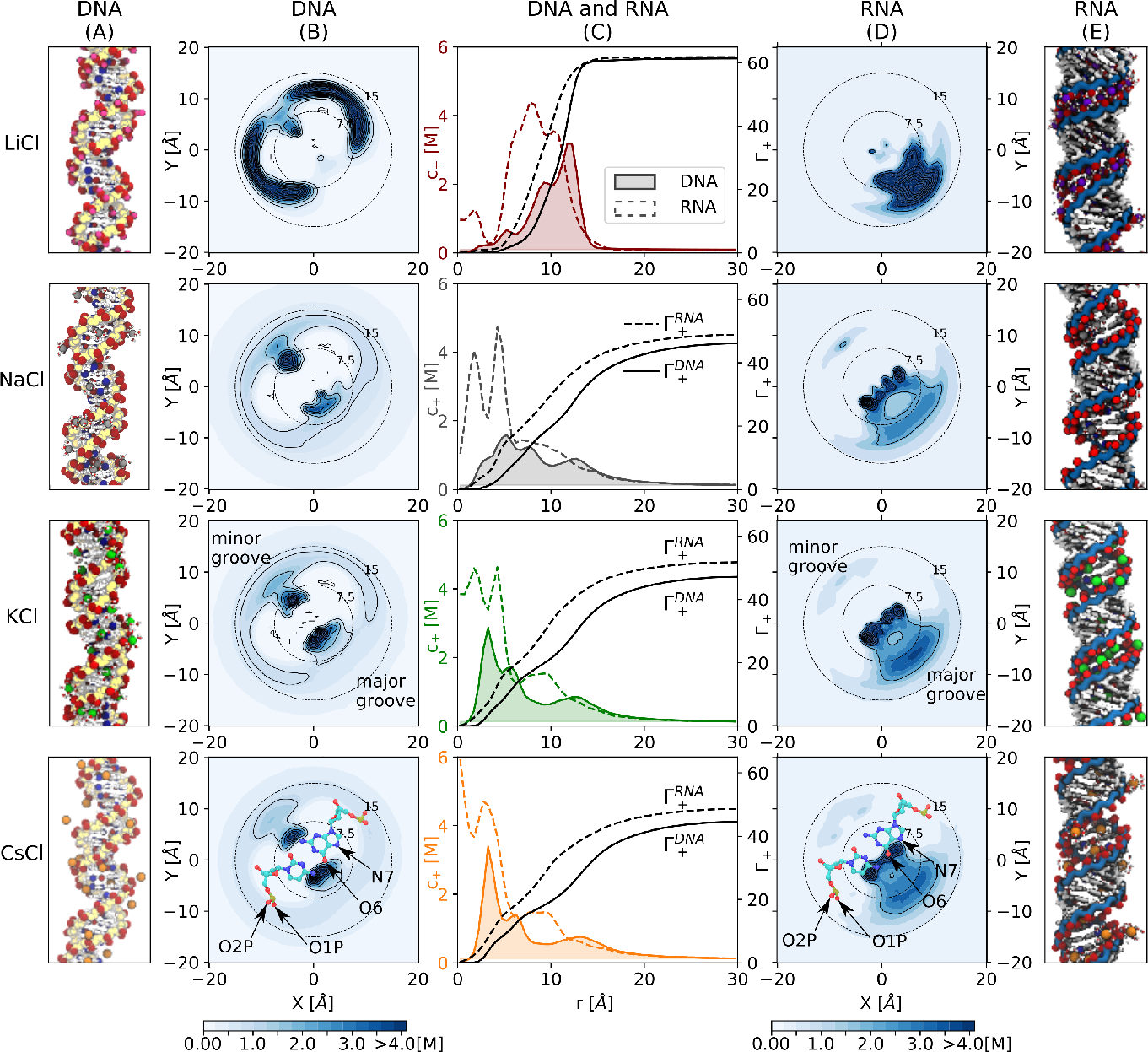
Ion distribution and excess of monovalent cations around DNA and RNA. (A,E) Simulation snapshots. Backbone atoms are indicated in yellow and blue for DNA and RNA, respectively. The most frequent ion binding sites are highlighted: red (O1P, O2P, and O6 atoms) and blue (N7 atoms). (B,D) Top view of the untwisted helicoidal ion concentration obtained with the software canion [79, 80]. In this representation, the upper-left and lower-right corners correspond to the minor and major grooves as indicated by the superimposed molecular schemes of cytosine-guanine (bottom). In these schemes the most frequent ion binding sites are labeled. The dotted concentric circles indicate the distance to the center of the helix (radius in Å). (C) Ion concentration profiles *c*_+_ as function of the distance *r* for DNA (solid line) and RNA (dashed line). Concentration profiles for DNA are filled for clarity. Figure reproduced from Ref. [77] with licence CC-BY 4.0.

The distributions of all the cations can be divided into two regimes: long and short distances from the surface of the DNA (Fig. 4C). Far away from the nucleic acid surface, the cylindrical cations concentration decay towards the bulk value, as predicted by classical mean-field theories [81, 82] (see also section 1). However, at short distances from the DNA helix (*r* 15 Å), the ion distributions are highly structured and unique for each ion type and nucleic acid type. The ion distributions close to the nucleic acid surface emerge from the direct interactions of the ions with the binding sites at the nucleic acids, as described in the previous section.

High and low charge density cations preferentially interact with the backbone and the nucleobase binding sites of nucleic acids, respectively. For example, all DNA profiles in Fig. 4 have a local maximum at the position of the backbone binding sites *r* ≈ 11 − 13 Å, i.e., all the cations interact at the backbone of DNA. In addition, this peak is prominently higher for ions with high charge density, e.g., Li^+^ compared to ions with intermediate or lower charge density, such as K^+^ or Cs^+^. In contrast, the innermost peaks (*r <* 10 Å), corresponding to the interaction with the nucleobases at the minor and major grooves (see Fig. 4B), increase with decreasing ion charge density (Li^+^ *<* Na^+^ *<* K^+^ *<* Cs^+^).

The simulations reveal that despite having helices with equal total charge and analogous sequences the ionic distributions depend both on ion type and the type of nucleic acid. Each cation type follows a distinct distribution when interacting with DNA or RNA. Simultaneously, for all ions, the ionic distribution around DNA is notably different from the distribution of the same ion around RNA. In DNA, which is normally a B-helix, ions are distributed between the major groove, the minor groove, and the backbone, whereas RNA, which typically is an A-helix, always favors ion localization in the major groove. As expected from the dinucleotide results (section 2.2.1), ions with high charge density interact strongly with the nucleic acid backbone, whereas ions with low density interact with nucleobases. The ion-site affinities explain the preferential distributions around DNA because in the B-helix the binding sites are spatially segregated. However, in the case of RNA, the interactions between the individual binding site and ions are insufficient to describe the results. We need to consider the topology of RNA. The A-helix points the backbone binding sites toward the interior of the major groove, where the nucleobase binding sites are also located.

In summary, we used MD simulations and recent development in atomic models [72, 73, 71, 65, 74] to quantify, in detail, the interaction of ions with nucleic acid binding sites [77, 9, 71]. The simulations showed high selectivity of ion binding to nucleic acids. Taken together, this detailed study allows us to understand a variety of experiments, to visualize how ions change the electrostatic environment around nucleic acids, and with it, the RNA properties and its interactions with other molecules.

## 3 RNA mechanical properties

The mechanical properties of nucleic acids play a key role for their functioning in vivo as they affect e.g. the affinity of binding proteins [83] or the packaging of genetic material in viral capsids. Experimental and simulation works have outlined a rich landscape for the elastic response of nucleic acids, unveiling a similar behavior of RNA and DNA in some properties, yet also showing marked differences in several other features. The current knowledge of nucleic-acids elasticity is mostly oriented towards duplexes. Several notable exceptions include experiments on single-stranded nucleic acids[84, 85, 86, 87, 88, 89, 90, 87, 91, 92, 93], which have highlighted a pronounced higher flexibility when compared to their double-stranded counterparts. Recent simulations [94, 95] and theory [90, 93, 96] have also provided fresh insights, although overall the state of the art is not as developed as for double-stranded nucleic acids, which will be the focus of the present section.

### 3.1 Local elasticity: base-pair-step parameters

A powerful insight on molecular flexibility can be obtained by the study of fluctuations, since more rigid molecules are characterized by weaker thermally-induced conformational fluctuations. Based on this, atomistic simulations demonstrated that for various angular degrees of freedom double-stranded RNA (dsRNA) is stiffer than double-stranded DNA (dsDNA), including e.g. the sugar pucker and the glycosidic angle[97]. Nevertheless, later simulation work has shown that the relative flexibility between dsDNA and dsRNA depends on the particular mechanical stress begin considered, as well as the scale at which it is being applied[98, 99]. In order to characterize the flexibility of nucleic acids, one has first to introduce suitable theoretical frameworks, which we rapidly describe in this section.

In complex molecules such as nucleic acids, the conformational state is usually characterized by introducing multiple variables which are significantly coupled with each other[99, 100]. Considering a perturbative approach, the elastic energy *U* is harmonic around the ground state[101]:

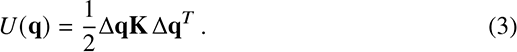

In the previous formula, **q** = {*q*_1_*, . . ., q_N_* } is a vector containing the *N* conformational variables; Δ**q** = **q**−**q**_0_, where **q**_0_ contains corresponds to the minimum-energy state; the apex *^T^* indicates transposition; and **K** is the stiffness matrix, where the diagonal elements are the elastic constants associated to the various modes and the off-diagonal elements are the coupling constants. Application of the equipartition theorem to this quadratic energy leads to

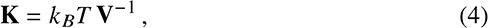

where *k _B_T* is the thermal energy and **V** is the covariance matrix, with elements *V_ij_* = Δ*_qi_*Δ*q_j_* and ⟨· · · ⟩ denoting thermal averaging. The previous formula provides a handy method to determine the stiffness matrix **K** based on the knowledge of **q**, acquired from different replica of the system or measuring the same thermally-equilibrated sample at different times.

A popular choice to characterize local elasticity of duplexes is provided by the base-pair-step parameters[101, 99], which give the relative position (shift, slide, rise) and orientation (tilt, roll, twist) between the planes of consecutive base pairs.

By setting **q** = {shift, slide, rise, tilt, roll, twist}, the local elasticity of dsDNA and dsRNA has been assessed by database analysis of crystal structures[101, 99] as well as by atomistic simulations[99]. This analysis has shown that, on average, dsRNA is more rigid than dsDNA at the local level. Yet, there is a wide variability according to sequence, which can be exploited to easily device dsRNA molecules which are actually softer than dsDNA with the same sequence. Moreover, even for the same sequence, the relative flexibility of dsRNA and dsDNA sometimes depends on the particular conformational feature being addressed. For instance, for the base-pair step CG, dsRNA is stiffer than dsDNA for the slide, but it is softer for the rise[99].

### 3.2 Global elasticity: elastic rod model

For global features, a nucleic acid is usually ascribed to a homogeneous, continuous rod characterized by stretch, twist, and bend deformations. For molecules up to tens of base pairs, the bending fluctuations of dsRNA and dsDNA can be neglected. Moreover, also for long molecules (as the ones considered in single-molecule experiments) the bending effects can be uncoupled from the other elastic modes when large pulling forces are considered [102]. By focusing on stretch and twist conformational changes, one thus chooses **q** = {*L, θ*} in Eq.(3), with *L* and *θ* being the overall contour length and torsion, respectively. In this case, Eq.(3) is usually written in the

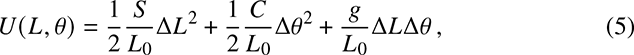

where *L*_0_ is the equilibrium value for the contour length, Δ*L* and Δ*θ* are the deviations for contour length and torsion, while the elastic constants *S, C* and *g* are the stretch modulus, the twist modulus and the twist-stretch coupling, respectively.

Eq.(5) characterizes the energy of the so-called elastic rod model. Based on Eq. (4) and Eq.(5), one can obtain the elastic constants from the fluctuations of *L* and *θ*. An alternative route consists in applying a force *f* and/or a torque *τ*, and minimize the total energy *U* − *f* Δ*L* − *τ*Δ*θ*. Thanks to the formulas obtained, fit of the curves giving ⟨Δ*L*⟩ and ⟨Δ*θ*⟩ as a function of *f* and *τ* enables computing the elastic constants.

This approach implicitly assumes that the elastic constants are independent of the applied mechanical stress.

The values of *S, C* and *g* obtained in single-molecule experiments are recapitulated in Table 1 top. While the twist response is similar for both duplexes, dsRNA is significantly softer than dsDNA for stretching. This is in contrast with the elasticity at the local level, where on average dsRNA was found to be stiffer, and provides a further example of the dependence of the relative flexibility between dsRNA and dsDNA on the deformation mode under consideration[99]. A remarkable qualitative difference between dsDNA and dsRNA emerges in the twist-stretch coupling, which has opposite sign for the two molecules [11]. This implies that, upon pulling, dsRNA relaxes its torsional state, while dsDNA overwinds. The sequence dependence of the elastic constants has hitherto not been experimentally characterized in detail, although it has been shown that dsDNA rich in A-tracts is generally stiffer than a random sequence [103]. Finally, it has been reported that for both dsRNA and dsDNA the stretching modulus increases with the ionic strength of the embedding solution, i.e. when the effect of electrostatics is weakened [12, 104].

**Table 1.**
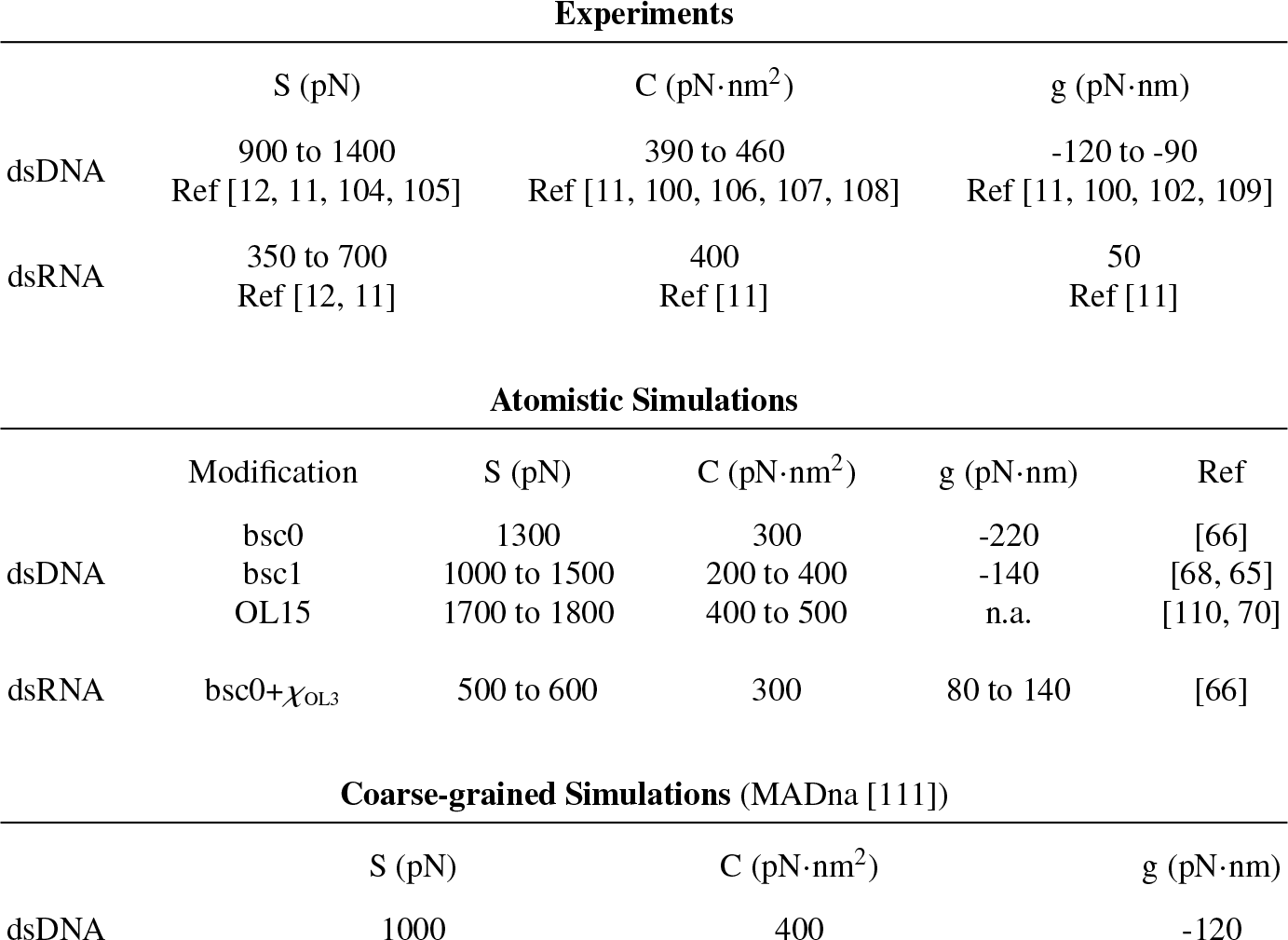
Collection of elastic constants obtained in the literature for dsDNA and dsRNA.

| Experiments |  |  |  |  |  |
| --- | --- | --- | --- | --- | --- |
|  | S (pN) | C (pN·nm<sup>2</sup>) | g (pN·nm) |  |  |
| dsDNA | 900 to 1400 | 390 to 460 | -120 to -90 |  |  |
|  | Ref [12, 11, 104, 105] | Ref [11, 100, 106, 107, 108] | Ref [11, 100, 102, 109] |  |  |
| dsRNA | 350 to 700 | 400 | 50 |  |  |
|  | Ref [12, 11] | Ref [11] | Ref [11] |  |  |
| Atomistic Simulations |  |  |  |  |  |
|  | Modification | S (pN) | C (pN·nm<sup>2</sup>) | g (pN·nm) | Ref |
| dsDNA | bsc0 | 1300 | 300 | -220 | [66] |
|  | bsc1 | 1000 to 1500 | 200 to 400 | -140 | [68, 65] |
|  | OL15 | 1700 to 1800 | 400 to 500 | n.a. | [110, 70] |
| dsRNA | bsc0+$\chi_{OL3}$ | 500 to 600 | 300 | 80 to 140 | [66] |
| Coarse-grained Simulations (MADna [111]) |  |  |  |  |  |
|  | S (pN) | C (pN·nm<sup>2</sup>) | g (pN·nm) |  |  |
| dsDNA | 1000 | 400 | -120 |  |  |

Atomistic simulations have provided a fundamental contribution to the microscopic understanding of nucleic-acids elasticity. Indeed, the experimental results for *S, C* and *g* were reproduced with good accuracy by all-atom simulations performed on short sequences (Table 1 center). The simulations captured quantitatively the stretching modulus and, although with large variability, the twist modulus. Moreover, they correctly account for the opposite sign of *g*. A microscopic analysis of the simulations enabled to ascribe this qualitative difference to the different sugar pucker induced by the presence or absence of the hydroxyl group in the ribose [66]. Quantitatively, the magnitude of *g* is overestimated by the simulations [66, 68], although this can be partially ascribed to finite-size effects introduced by the short length of the simulated molecules (∼20 base pairs) [111].

In order to simulate nucleic acids at larger scales, due to computational limits one needs to introduce coarse-grained descriptions. In this regard, until recently the mechanics of dsDNA was not satisfactorily captured by existing models, either due to the lack of sequence dependence or to the overestimation of the elastic constants [111, 112, 113]. Moreover, the sign of *g* was not captured, indicating inherent limits in the qualitative assessment of dsDNA mechanics. The accuracy of atomistic simulations has recently enabled the development of MADna, a sequence-dependent model focused on the conformational and mechanical properties of dsDNA, and entirely parameterized from atomistic simulations [111]. Despite its coarse-grained nature, MADna quantitatively reproduces the main sequence-dependent conformational and elastic properties obtained in atomistic simulations and experiments (see Table 1 bottom for average elastic constants). Moreover, it also accurately captures the sequence-dependent helical pitch as obtained from cyclization experiments [114], as well as the pronounced stiffness of A-tracts [103]. The development of coarsegrained models for the elasticity of dsRNA is still in its infancy, in comparison to dsDNA. At present, there is only a report of the stretch modulus for one such model, showing that *S* is significantly underestimated [115]. Given the importance of nucleic-acids elasticty, there is clearly a great need for further development of dsRNA models.

### 3.3 Global elasticity: persistence length

For longer molecules, also thermally-induced bending has to be taken into account. In this regard, it is customary to resort to the tools provided by polymer theory. The self-correlation of the tangent vectors *t*^ along the chain is predicted to decay exponentially with the contour length Δ*s* separating the points under inspection [116]: 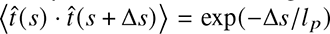, where the persistence length *l _p_* gives the length scale of thermally-induced bending fluctuations. Experimental imaging via Atomic-Force Microscopy can be used to determine the correlation function, from which the value of *l _p_* can be extracted [117]. Moreover, the wormlike chain model provides a handy formula for the response of the chain to a pulling force *f* [118]. Based on either of these approaches, experiments showed that *l _p_* = 45 − 55 nm for dsDNA [12, 11, 104, 105, 117, 110], while *l _p_* = 57 − 63 nm for dsRNA [12, 11, 119].

Therefore, dsRNA is stiffer than dsDNA when it comes to global bending. Moreover, at low pulling forces (*f <* 1 pN) the coupling between bending and twist results in an effective smaller twist modulus [120, 121], while for both dsDNA and dsRNA the persistence length decreases with the ionic strength of the solution [12, 104, 105].

Finally, cyclization experiments on short dsDNA fragments of repeating patterns investigated the sequence dependence of the persistence length [114].

Atomistic simulations for dsDNA based on AMBER force fields account with good accuracy for the experimental results. Particularly, it was found that *l _p_* = 43−51 nm for parm99+bsc0 [122], *l _p_* = 40 − 60 nm for parm99+bsc1 [65, 110, 123] and *l _p_* ≃ 60 nm for parm99+OL15 [110]. In the case of dsRNA, it was estimated that *l _p_* = 70 − 80 nm for bsc0 [99] and bsc0+*χ*_OL3_ [123]. Hence, all-atom simulations capture the relative bending flexibility of dsDNA and dsRNA, although slightly overestimating the quantitative estimations.

At the coarse-grained level, several models give a correct estimation of *l _p_* for dsDNA[111, 112, 113, 124, 125, 94, 126], yet in various cases the parameters of the models were tuned for its optimization[112, 113, 124]. Notably, the TIS and 3SPN models capture the decrease of *l _p_* with the ionic strength [94, 112]. As for the sequence dependence, the best performance has been shown for MADna and cgDNA[111, 126], which accurately account for the values of *l _p_* obtained by cyclization experiments[114]. Finally, both MADna and oxDNA capture the effect of twist-bend coupling [111, 127]. In the case of dsRNA, there are only few works estimating *l _p_* for coarse-grained models. Its value is usually underestimated [115, 128, 129], although a suitable parameterization of the MARTINI model can reproduce the correct result [128].

### 3.4 Force dependence of elastic constants

Experimental evidence has pointed out that the elastic constants of nucleic acids depend on the pulling force *f* . For instance, stretched dsDNA overwinds until *f* ≃ 35 pN (which implies *g <* 0), while it underwinds for larger forces (*g >* 0)[100, 102]. On this matter, atomistic simulations have provided unique fundamental insight.

However, a theoretical challenge first needed to be overcome, since neither of the approaches presented above is suitable to study stress-dependent elasticity: fitting full stress-dependent curves impedes by construction to extract the stress dependence of constants, while the formula for fluctuations (Eq. (4)) is valid only for harmonic energies, which are perturbed by the linear terms in which the force or torque appear. In this regard, at a general level the energy can be written as [130]

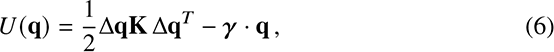

where γ is a vector containing the mechanical stresses associated to the various deformation modes. For instance, for **q** = {*L, θ*} one has γ = { *f, τ*}, where *f* is the pulling force and *τ* the applied torque. Application of the generalized equipartition theorem gives [130]

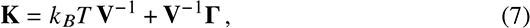

where Γ*_ij_* = ⟨Δ*q_i_*⟩ *γ _j_*. This approach was employed [130] to analyze the atomistic simulations from Refs.[66, 131, 132], where various dsDNA and dsRNA molecules with heterogeneous sequences were pulled with forces up to 20 pN. It was found that the stretching modulus of dsDNA increases with force, while that of dsRNA decreases. This unexpected result was rationalized by observing the opposite change in slide upon pulling the two molecules, which induces a strengthening and weakening of stacking interactions for dsDNA and dsRNA, respectively [130]. In contrast, the twist modulus was found to decrease for both kinds of duplexes. This behavior was ascribed to the progressive alignment of the normals to the base planes with the helical axis at increasing forces, which eventually identifies local torsion with the helical twist. A toy model based on this conjecture was capable of quantitatively reproducing the results for dsDNA without any fitting parameters, supporting this physical picture [130]. Finally, it was also found that, when considered with sign, *g* increases with force. Since for dsDNA *g <* 0 at low forces, this is in line with the experimental observations that for strong enough pulling the sign is reverted. All in all, these results add a further layer of complexity to the already rich landscape of nucleic-acids elasticity, where microscopic changes in the chemical structure result in strikingly different macroscopic mechanic behavior.

## 4 RNA genome 3D reconstruction

The assembly process in RNA viruses is a complex phenomenon which involves a cooperative interplay between the genome and the capsid proteins [1]. Consequently, the packaged structure of RNA in the virus’ interior is of fundamental importance for its stability, and can shed light into the details of the assembly or disassembly processes[133]. As we saw in the sections above, mean-field theoretical approaches have pointed out the effect of persistence length, secondary structure, branching degree and electric charge into the assembly/disassembly of the packaged structure [134, 135, 136, 137]. Other authors have stressed the importance of sequence-specific interactions between the genome and the capsid proteins, whose identification can lead to a sequential description of the assembly process[133]. A systematic and concrete picture of the geometrical conformation of the genome inside the virus can complement these results and be used for generating practical, all-atom intraviral RNA conformations. In fact, many experimentally determined structures contain some RNA fragments resolved, which have been used for proposing 3D models [138]. Such is the case of Satellite Tobacco Mosaic Virus (STMV) [78], Bacteriophage MS2 virus [139] and Cowpea Chlorotic Mottle Virus [140], among others. The case of STMV is of particular interest, since its crystal structure displays contains 30 double helices which comprise around 60% of its genome, although it lacks specificity about its sequence. It is therefore a good starting point for testing the 3D reconstruction in two aspects: a full model based in the secondary structure, and another one which considers the geometrical restraints imposed by the experimental crystallographic data of the STMV [27]. Given the size of the RNA involved, it is recommended to work, in an initial phase, with simpler descriptions. Coarse-grained models [69] offer a good trade-off between accuracy and efficiency, and can help to distinguish realistic models from spurious assemblies with clashes or unfeasible topological artifacts.

### 4.1 Fragment assembly

Several well-established approaches generate three-dimensional RNA models from the assembly of experimentally determined fragments. Knowing the secondary structure, radius of gyration or tertiary contacts greatly helps to distinguish the most suitable structures among the large number of possibilities given a specific scoring function [141, 142, 143, 144, 145, 146]. Inside the virus, the RNA might adopt non-trivial conformations mainly due to the confinement and the interaction with the capsid proteins, so the use of experimentally determined fragments could be too restrictive. We have then generated fragments of the secondary structure elements, hairpins, stems, and junctions, through on-the-fly short coarse-grained simulations using secondary structure restraints[27, 58]. Figure 5 shows an example on a fragment of STMV. Stems and hairpins are generated and later assembled with their closest junction, which is sampled from a simulation where the energy function contains only secondary structure and radius of gyration restraints, excluded volume and backbone connectivity. The conformation of the junctions can be sampled by performing several independent short-simulations, in order to obtain a wide number of different geometries. After the assembly process, it is likely that the proximity of the assembled fragments will produce clashes between coarse-grained beads or, even worse, entanglements between the loops, stems, and junctions. These issues are circumvented by energy minimization and identifying and removing the links between the loops. The detection of these artifacts can be performed by known methods[147], and their removal can be done by applying repulsive energy terms on virtual sites situated at specific points of the loops, as illustrated in Figure 5 d)[27, 148].

**Fig. 5.**
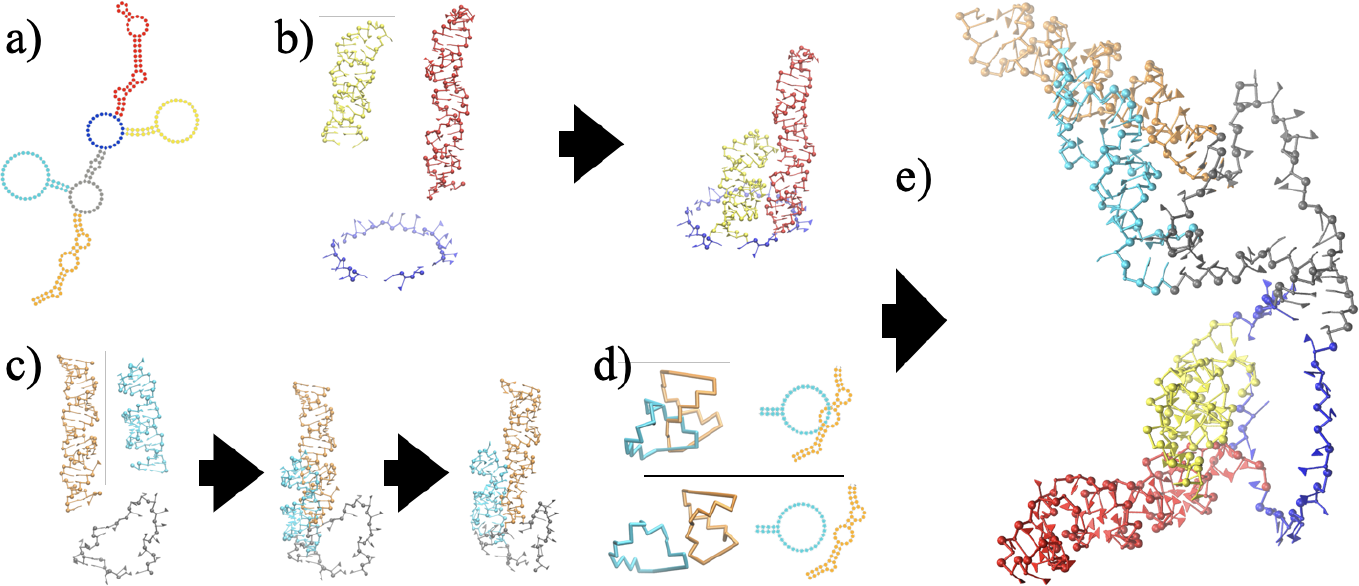
Assembly process illustrated for nucleotides 330 to 575 of STMV, a) in virio secondary structure. b)Two loops containing hairpins are folded separately and attached to a junction, generated independently. c) The procedure is repeated in another junction. In this case, the assembled fragments exhibit clashes and are entangled in a non-trivial manner. d) Zoom of the entanglement, in backbone (left) and secondary structure (right) representation: each fragment forms a ring by joining the phosphate and sugar groups, closed by canonical base pairs. The rings are represented in the unrefined (top) and refined (bottom) conformations. e) The whole structure is assembled, following energy minimization and removal of clashes and links between secondary structure elements [27].

In a later step, the generated structures can be submitted to external forces which confine them to a spherical domain. Nevertheless, the structure might have additional restraints coming from the experimental data and must be treated in a specific way.

### 4.2 eRMSD restraints

The imposition of geometric restraints requires a parameter to compare the structural differences between a given structure and a template. The eRMSD distance [149] is our choice since it is a specific metric for dealing with nucleic acids, taking into account the difference of the spatial arrangements of the nucleobases belonging to two structures. For this aim, it defines a local reference frame on each nucleobase, which is situated at the center of mass of their C2, C4 and C6 atoms. For each pair of bases *i* and *j*, the vector joining their centroids in the local reference frame of nucleobase *i* is denoted by **r***_ij_*, which is rescaled in order to take into account the nucleobase shape, by defining **r̃***_ij_* = (*r_x_*/*a, r_y_*/*a, r_z_*/*b*), with *a* = 5Å and *b* = 3Å. The eRMSD between two structures *α* and *β* composed by *N* nucleotides is given by

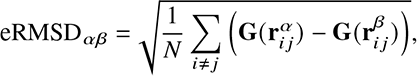

where **G**(*r*) is a four-dimensional vector which satisfies |**G**(**r̃***^α^*) − **G**(**r̃***^β^*)| ≈ |**r̃***^α^* − **r̃***^β^* | for **r̃***^α^,* **r̃***^β^* ≪ *D_c_* and |**G**(**r̃***^α^*) − **G**(**r̃***^β^*)| = 0 for **r̃***^α^,* **r̃***^β^ > D_c_*, with *D_c_* a dimensionless cutoff. The eRMSD restraint is defined as a harmonic potential energy term of the form

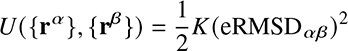

where *K* is a constant with units of energy. The energy is imposed on a set of nucleotides with respect to its template, which can also be used for imposing secondary structure restraints. In our case, we have tested it on a set of double helices defined from the secondary structure proposed by Weeks and co-workers [63]. We took nucleotides 772 to 897, as illustrated in Figure 6a), and chose five stems around a five-fold symmetry axis from PDB 4OQ9 (Figure 6 b)) for defining two templates: one where the hairpins point outwards (case A, Figure 6c)) and another where they point inwards (case B, Figure 6d)). An initial condition can be easily generated using the procedure described in the previous section. The eRMSD restraints can be applied on any representation of RNA which is able to define the orientation of the nucleobases. In the present case, we have employed the SPQR coarse-grained model [58], which has this feature incorporated into its code. The model represents each nucleotide by a triangular nucleobase and two beads for the sugar and phosphate group.

**Fig. 6.**
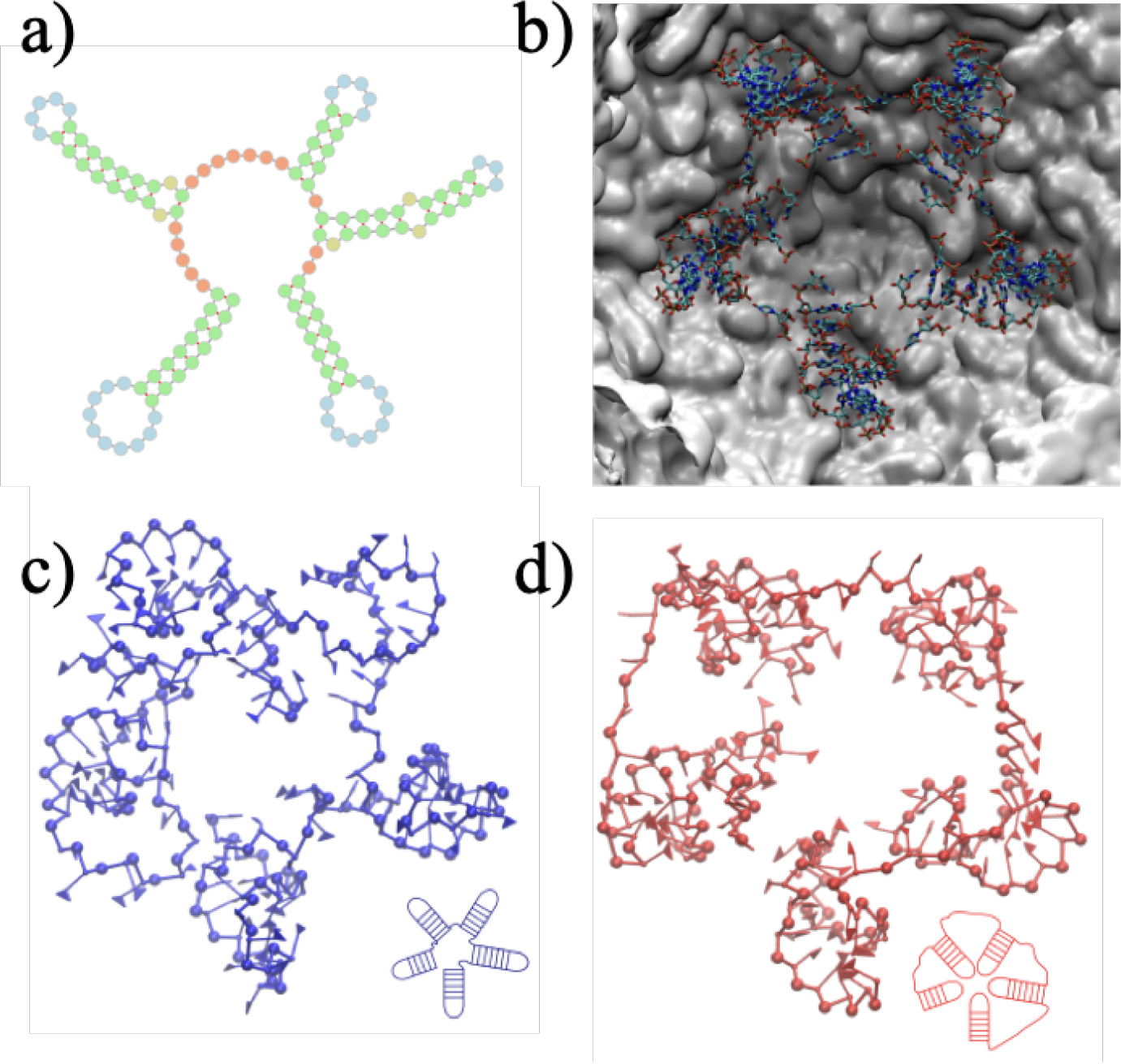
Five consecutive hairpin loops are mapped according to different templates. a) Secondary structure of the fragment. b) Template taken from PDB structure. c) Case A, after simulation d) Case B, after simulation. Figure reproduced from [27] with licence CC-BY 4.0.

The value of the eRMSD and backbone energy are good descriptors of the suitability of the proposed configurations, as shown in Figure 7. After running 3 short independent Monte Carlo simulations for each case, we see that these parameters can blindly distinguish that case A is more stable and favorable than case B, and therefore, selected for backmapping into atomistic detail. The procedure could be iterated over the rest of the structure in order to obtain a reduced set of possible conformations of the whole arrangement of the STMV genome inside the virus.

**Fig. 7.**
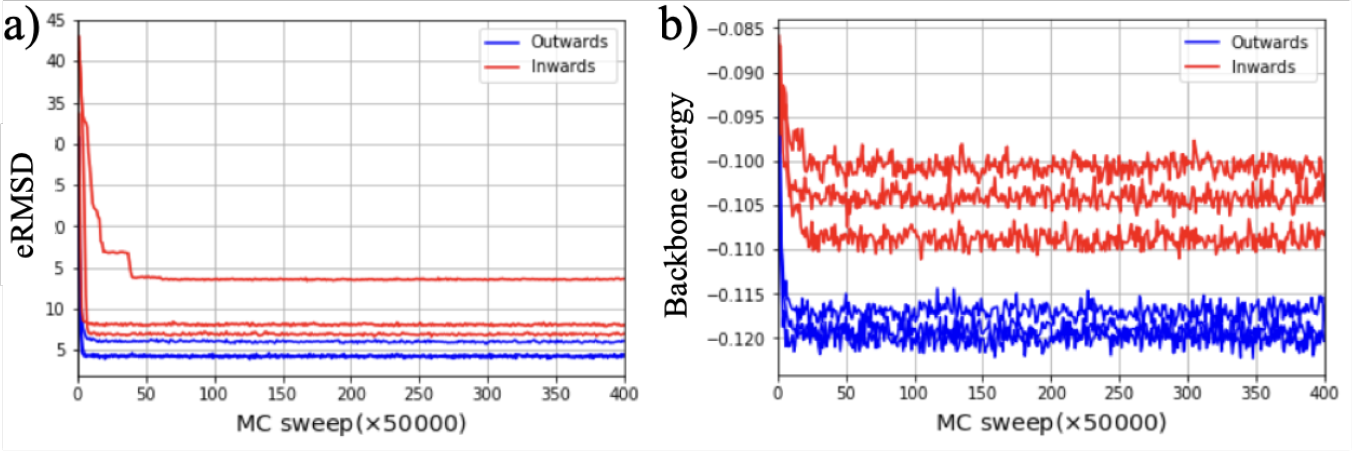
a) eRMSD between the 5 stems and their templates. b) Backbone energy for both conformations as a function of MC sweep. Figure reproduced from [27] with licence CC-BY 4.0.

## 5 Discussion and Prospective methods

The studies described in the previous sections outline the exquisite multiscale nature of RNA physics. The assembly of a viral capsid, as well as a virion requires to consider electrostatic interactions, determining feasible conformations of the genome and their effects on the nanomechanical stability of the capsid and genome itself. Here, we have shown the potential of atomistic, coarse-grained simulations and continuum models for modeling RNA and its interactions at different spatial and temporal scales. Still, there are several challenges ahead for RNA physics virology. Determining the folding of RNA and its dynamics within the virus remains a challenging open problem.Experimental techniques such as NMR, CryoEM, or recently CryoET have had dramatic progress recently and now allow us to visualize the nanoscopic details of the interior of the virions [150]. However, the ’dynamic structure’ of RNA is commonly difficult to efficiently sample by pure experiments[151, 152]. Hence, high-resolution experiments are complemented with computational models. For example, computational approaches were used to refine 3D RNA reconstructions with additional restraints on tertiary contacts given by SAXS experiments [153, 148]. Furthermore, based on chemical probing and CryoEM, it was possible to reconstruct the frameshift stimulating element from SARS-CoV-2 all the way down to atomistic resolution [24]. Likewise, a general solution remains out of reach. However, after the astonishing success of Alphafold for protein folding [154] and similar developments for RNA [155, 25], machine learning-based methods hold a great promise for solving RNA folding.

Realistic models of whole virus dynamics require physics-informed models. Yet, a current problem with multiscale approaches is that they remain largely disconnected. No integrative model takes also into account detailed electrostatics and mechanical properties of both RNA and the proteins forming the non-enveloped viruses. The mechanics of RNA is vital for its packaging in a confined medium such as a viral capsid, yet scalable computational approaches focused on this feature are still at their infancy.

Similarly, electrostatics is central for the assembly, disassembly and stability of virus capsids [2], and virion assembly [3]. However, it remains unclear which is the required precision to describe the interactions between RNA and capsid-proteins, ions and short range interactions. Recent research has been used free energy profiles extracted from atomic calculations as effective interactions in coarse-grained models [156] and also multiscale [8]. Similar approaches are already under discussion [1], which could include important interactions towards single amino-acids in proteins [18], water or surfaces into multiscale simulations [157]. In any case, there is little doubt that the exceptional challenge posed by RNA organization within viruses will be overcome only via a synergistic approach complementing computational approaches at multiple scales with the growing amount of experimental knowledge. The integrative approach is complex itself, nonetheless, current developments to discern between molecular interactions can be used to test physical assumptions, guide the interpretation of experiments and possibly the direction of new experiments. Finally, a robust parametrization of the underlying RNA-protein molecular interactions (mechanical and electrostatics) would consequently permit the modification and/or control the capsid status from assembly to disassembly or vice versa, which are awaited features for the next generation Virus-like Particles (VLPs) nanomedicine applications.

## Acknowledgements

H.V.G. acknowledges the core funding support from the Slovenian Research Agency, under grant No. P1-0055, and the financial support of the Community of Madrid and the European Union through the European Regional Development Fund (ERDF), financed as part of the Union response to Covid-19 pandemic. S.A. received the support of a fellowship from “la Caixa”Foundation (ID 100010434) and from the European Union’s Horizon research and innovation programme under the Marie Sklodowska-Curie grant agreement No. 847648. The fellowship code is LCF/BQ/PI20/11760019. S.A. acknowledges support from the Ministerio de Ciencia e Innovación (MICINN) through the project PID2020-115864RB-I00 and the “María de Maeztu” Programme for Units of Excellence in R&D (grant No. CEX2018-000805-M). S. P. acknowledges the project FONDECYT Iniciación en Investigación 11181334 for financial support.

## Footnotes

1 Details on the free energy profiles calculations can be found in Ref. [9]

